# Modular Dynamic Biomolecular Modelling with Bond Graphs: The Unification of Stoichiometry, Thermodynamics, Kinetics and Data

**DOI:** 10.1101/2021.03.24.436792

**Authors:** Peter J. Gawthrop, Michael Pan, Edmund J. Crampin

## Abstract

Renewed interest in dynamic simulation models of biomolecular systems has arisen from advances in genome-wide measurement and applications of such models in biotechnology and synthetic biology. In particular, genome-scale models of cellular metabolism beyond the steady state are required in order to represent transient and dynamic regulatory properties of the system. Development of such whole-cell models requires new modelling approaches. Here we propose the energy-based bond graph methodology, which integrates stoichiometric models with thermodynamic principles and kinetic modelling. We demonstrate how the bond graph approach intrinsically enforces thermodynamic constraints, provides a modular approach to modelling, and gives a basis for estimation of model parameters leading to dynamic models of biomolecular systems. The approach is illustrated using a well-established stoichiometric model of *Escherichia coli* (*E. coli*) and published experimental data.

## 1 Introduction

The recent explosion of omics data has generated an interest in developing dynamic whole-cell models that account for the function of every gene and biomolecule over time. Such models have the potential to “predict phenotype from genotype” [1–3] and hence to “transform bioscience and medicine” [4]. Critical to understanding the large-scale metabolism within cells is the stoichiometric approach [5– 8], which has had notable successes including the genome-scale reconstruction of the metabolism of *Escherichia coli* (*E. coli*) [9–11] and Neocallimastigomycota Fungus [12].

The stoichiometric approach can give rise to constraint-based models such as Flux Balance Analysis (FBA)[13], which predict metabolic fluxes at steady state. However, most implementations of such constraint-based models do not explicitly consider energy. This can lead to mass flows that are not thermodynamically possible because they violate the second law of thermodynamics. Such non-physical flows can be detected and eliminated by adding additional thermodynamic constraints, as in *Thermodynamics-based Metabolic Flux Analysis* (TFA)[14, 15], *Energy Balance Analysis* (EBA) and *Expression, Thermodynamics-enabled FLux models* (ETFL) [16–20] and loopless FBA [21]. Whereas constraint-based models provide metabolic fluxes, they generally do not explicitly account for metabolite concentrations, or how fluxes vary over time, both of which are required for dynamic whole-cell modelling. However, the stoichiometric approach can help to bridge towards dynamic models capable of satisfying these requirements. In this context, there has been work into developing two types of large-scale dynamic models: fully detailed mass action stoichiometric simulation (MASS) models [7, 22, 23] and simplified network models that use non-mass action rate laws such as lin-log laws or modular rate laws [24, 25]. Although mass-action approaches seem restrictive, we note that models of enzyme kinetics can be built from elementary mass-action reactions [26].

MASS models are parameterised by reaction rate constants which are subject to thermodynamic constraints such as the Wegscheider conditions [27] (Wegscheider conditions are a formulation of *detailed balance* conditions which avoid models which are inconsistent with thermodynamic laws [26, § 1.5]). This paper focuses on the mass-action formulation and introduces an alternative to MASS which explicitly incorporates thermodynamics. Specifically, the approach uses an alternative parameterisation related to that of Thermodynamic-Kinetic Modelling (TKM) [27, 28]. TKM explicitly divides parameters into those associated with capacities and resistances by analogy with electrical systems; this approach gives thermodynamic consistency without invoking additional constraints such as the Wegscheider conditions [27, 28]. Mason and Covert[29] developed a similar approach for a non-mass-action rate law.

Recently, the bond graph approach from engineering [30–33] has been adapted to biochemistry [34–37]. Bond graphs are close in spirit and application to TKM in that they produce ordinary differential equations for dynamic simulation [37] and that their parameters satisfy thermodynamic consistency without the need to invoke Wegscheider conditions [34–37]. However, bond graph models are endowed with several additional features:

1. Bond graphs can be easily generalised to model multi-physics systems and thus readily incorporate the physics of electrically charged species into an integrated model combining both chemical and electrical potential [38–42].
2. Bond graphs are modular [43, 44], a key requirement of any large-scale modelling endeavour [45].
3. Bond graph models can be systematically modified to give simpler bond graph models which remain compatible with thermodynamic laws [37, 46, 47].

The stoichiometric matrix of a biomolecular network can be derived from the corresponding bond graph [37, 43]. Similarly, as shown herein, a bond graph model can be constructed from a stoichiometric matrix. Thus, the large repository of models of biomolecular systems available in stoichiometric form are available as templates for developing bond graph models; we provide a methodology for this later in the paper. Furthermore, once rate laws such as mass action are added, such templates provide a basis for complete dynamic models of metabolic systems.

A key challenge in the development of dynamic models is the fitting of parameters to experimental data, especially when thermodynamic constraints need to be satisfied [48, 49]. For large-scale biomolecular models such as whole cell models, applying these constraints is particularly challenging [50]. In this paper, we use the thermodynamically safe parameterisation provided by bond graphs to resolve this issue. As in the TKM [27, 28] approach, the bond graph approach uses an alternative parameterisation which satisfies thermodynamic constraints *as long as the parameters are positive*; such inequality constraints are easier to handle than non-linear constraints. We illustrate this approach by generating a dynamic bond graph model of *E. coli* metabolism, using a well-established stoichiometric model [51] as a template and show that the use of thermodynamic parameters can significantly streamline the process of parameter estimation.

In summary, this paper proposes the fusion of the stoichiometric and bond graph approaches to modelling biological systems and illustrates its potential for the unification of stoichiometry, thermodynamics, kinetics and data.

§ 2 summarises the bond graph background to the rest of the paper. § 3 shows how bond graph models can be extracted from stoichiometric information, used to create modular models and analysed in terms of pathways; the relationship of the approach to Energy Balance Analysis is also discussed. § 4 applies these concepts to two subsystems within the *E. coli* core model – a well-documented [8, 51] and readily-available [52] stoichiometric model of a biomolecular system. § 5 shows how thermodynamically-consistent bond graph parameters can be extracted from experimental data and gives a dynamic simulation of the parameterised model. § 6 concludes the paper and gives directions for future work.

## 2 Bond Graphs

This section gives a brief introduction to the bond graph approach to modelling biomolecular systems based on the seminal work of Oster et al. [34, 35] as extended by Gawthrop and Crampin [37,44, 53].

### 2.1 Basic components

Bond graphs represent the energetic connections between components of a system. The *⇁* symbol is used to indicate an energetic connection, or ‘bond’, between components; the half-arrow indicates the direction corresponding to positive energy flow. In the biomolecular context, each bond is associated with two covariables: chemical potential *µ* (J mol^*−*1^) and flow *v* (mol s^*−*1^). The key point is that the product of *µ* and *v* is power *p* = *µv* (W). This ensures that models are consistent with the laws of thermodynamics, as energy flow is explicitly accounted for. In the context of cellular metabolism, and in line with the measurement of redox potentials, it is convenient to scale these co-variables by *Faraday’s constant F ≈* 96 485 C mol^*−*1^ to give

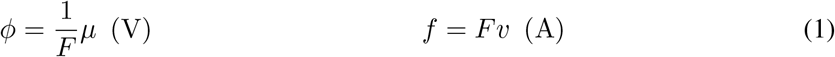

where (J C^*−*1^) has been replaced by the more convenient unit volt (V) and (C s^*−*1^) has been replaced by the more convenient unit ampere (A) [38]. As a useful rule-of-thumb, *µ* (kJ mol^*−*1^) can be converted to *ϕ* (mV) by dividing by 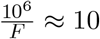.

Bonds transmit, but do not store or dissipate energy. Within this context, the bonds connect four distinct types of component:

#### 0 & 1 Junctions

provide a method of connecting two or more bonds, and therefore creating a network. Analogous to electrical systems, there are two types of junction, denoted **0** and **1**. The bonds impinging on a **0** junction share a common effort (chemical potential); the bonds impinging on a **1** junction share a common flow. Both **0** and **1** junctions transmit, but do not store or dissipate energy. As discussed previously [37], the arrangement of bonds and junctions represents the stoichiometry of the corresponding biomolecular system and thus the relationship both between reaction and species flows and between species potentials and reaction forward and reverse potentials. Furthermore, the reverse is also true: the stoichiometric matrix of a biomolecular system uniquely determines the bond graph, as will be discussed further below.

**Ce** represents biochemical *species*. Thus species A is represented by **Ce**:**A** with the equations:

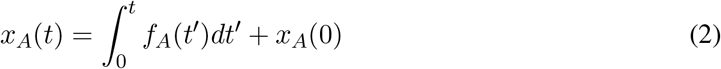

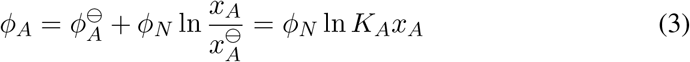

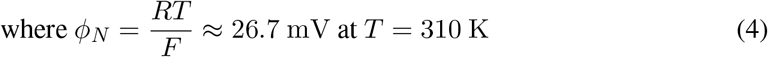

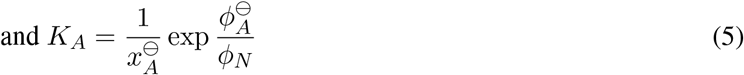

Equation (2) accumulates the flow *f*_*A*_ of species A. Equation (3) generates chemical potential *ϕ*_*A*_ in terms of the reference potential 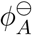 at reference conditions 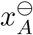. **Ce** components thus store, but do not dissipate, energy. An equivalent parameterisation that we use in this paper is to express the chemical potential in terms of *ϕ*_*N*_ and the species constant *K*_*A*_, as defined in Equation (5).

**Re** represents *reactions*. The flow *f* associated with each reaction is given by the *Marcelin – de Donder* formula [37, 54]:

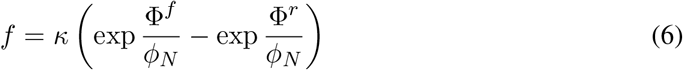

where Φ^*f*^ and Φ^*r*^ are the forward and reverse reaction potentials (or affinities), defined as the sums of the chemical potentials of the reactants and products respectively. If *κ* is constant, this represents the mass-action formula.

In general, *κ* is a function of Φ^*f*^, Φ^*r*^ and enzyme concentration [37]; for example, a reversible Michaelis-Menten formulation used in Gawthrop et al. [43] is:

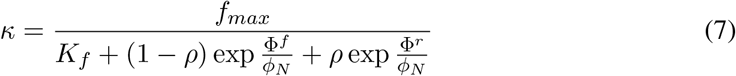

where the three constants *f*_*max*_, *K*_*f*_ and *ρ* define the kinetics. As discussed elsewhere [26, 37], enzyme kinetics can be modelled using the pair of reactions with mass-action kinetics

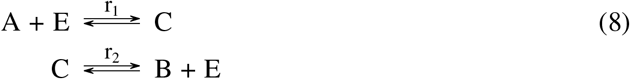

where A, B, E and C are the substrate, product, enzyme and complex of substrate and enzyme respectively; the bond graph representation is given in Appendix E. Equation (7) arises from the steady-state analysis of this model [37]. In particular:

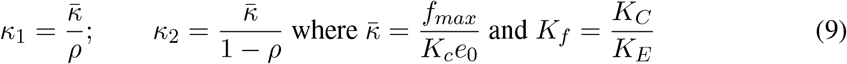

*K*_*C*_ and *K*_*E*_ correspond to Equation (5) for the complex and enzyme respectively and *e*_0_ is the total amount of enzyme (unbound and bound within the complex).

**Re** components dissipate, but do not store, energy. In general

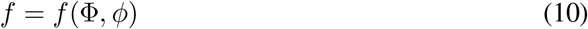

where Φ = Φ^*f*^ − Φ^*r*^ and *ϕ* is a vector containing the chemical potentials of every species. Since *f* always has the same sign as Φ, *f* () is dissipative in Φ for all *ϕ*:

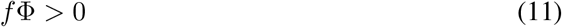

The key *stoichiometric* equations arising from bond graph analysis are [37]:

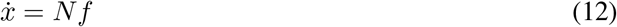

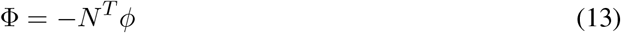

where *x, f*, Φ and *ϕ* are the species amounts, reaction fluxes, reaction potentials and species potentials respectively, all represented as vector quantities. *N* is the stoichiometric matrix of the network. Combining Equations (12) and (13):

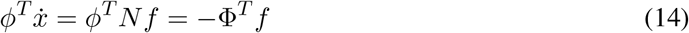

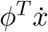 is the rate of energy into the species (which must be negative or zero for closed systems) and Φ^*T*^ *f* is the rate of energy dissipated by the reactions. Since 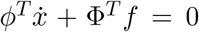, it follows that the network of bonds and junctions transmits, but does not dissipate or store, energy [37].

Moreover, the stoichiometric matrix *N* can be decomposed as [37]:

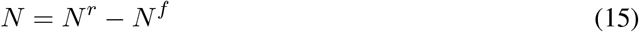

where *N*^*r*^ corresponds to the positive entries of *N* and *N*^*f*^ to the negative entries. The forward and reverse reaction potentials Φ^*f*^ and Φ^*r*^ are given by:

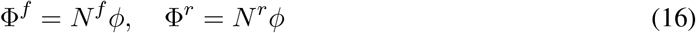

In other words, the stoichiometric matrix *N* can be derived from the system bond graph. § 3 shows that, conversely, the system bond graph can be derived from the stoichiometric matrix *N*.

### 2.2 Chemostats, Flowstats and Pathways

Modularity implies the interconnection of subsystems; thus such subsystems must be thermodynamically open. As discussed previously [38, 44], the notion of a *chemostat* [55] is useful in creating an open system from a closed system. The chemostat has a number of interpretations [38]:

1. one or more species are fixed to give a constant concentration [43]; this implies that an appropriate external flow is applied to balance the internal flow of the species.
2. as a **Ce** component with a fixed state.
3. as an external *port* of a module which allows connection to other modules.

In the context of stoichiometric analysis, the chemostat concept provides a flexible alternative to the primary and currency exchange reactions [6, 8, 56].

Alternatively, reaction flows can be fixed using the dual concept of *flowstats* [44], which has a number of interpretations:

1. as an **Re** component with a fixed flow.
2. as an external *port* of a module which allows connection to other modules.

In the context of this paper, we use flowstats to isolate parts of a network by setting the flows of certain reactions to zero. Such zero flow flowstats can also be interpreted as removing the corresponding enzyme via gene knockout.

In terms of stoichiometric analysis, the closed system equations (12) and (13) are replaced by:

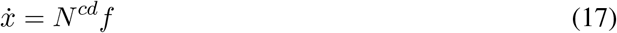

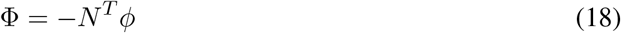

where *N*^*cd*^ is created from the stoichiometric matrix *N* by setting *rows* corresponding to chemostats species and *columns* corresponding to flowstatted reactions to zero [44]. As discussed by Gawthrop and Crampin [44], system pathways corresponding to Equation (17) are defined by the right-null space of *N*^*cd*^, that is, the columns of a matrix *K*_*p*_ satisfying the equation *N*^*cd*^*K*_*p*_ = 0. At steady state, the flows through these pathways are defined by:

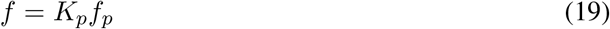

where *f*_*p*_ is the pathway flow. It follows from Equation (17) that Equation (19) implies that 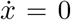. The *pathway* stoichiometric matrix *N*_*p*_ is defined as [53]:

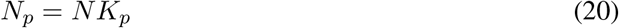

In a similar fashion to Equation (18), the pathway reaction potentials Φ_*p*_ are given by

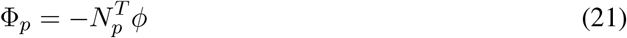

In the same way as the stoichiometric matrix *N* relates reaction flows to species and thus represents a set of reactions, the pathway stoichiometric matrix *N*_*p*_ also represents a set of reactions: these reactions will be called the *pathway reactions*.

Pathways can be divided into three mutually exclusive types [56] according to the species corresponding to the non zero elements in the relevant column of the *pathway* stoichiometric matrix *N*_*p*_:

**Type I** The species include primary metabolites; these pathways are of functional interest.

**Type II** The species include currency metabolites only; these pathways dissipate energy without creating or consuming primary metabolites. Such pathways are sometimes called *futile cycles*; however they have an important role to play in regulating metabolite flow [57–62].

**Type III** There are no species. These may arise when the same reaction is catalysed by different isoforms of the same enzyme.

Pathway reactions for type I pathways contain both primary and currency metabolites; pathway reactions for type II pathways contain currency metabolites only; pathway reactions for type III pathways are empty. The concept of pathways is applied to a simple example in Appendix B and to a biomolecular example (the Pentose Phosphate Pathway) in § 4.1.

## 3 Bond Graphs Integrate Stoichiometry and Energy

As discussed in the previous section, the stoichiometric matrix can be directly derived from the bond graph; this section shows that the converse is true and thus bond graphs can be automatically derived from preexisting stoichiometric representations thereby allowing bond graph energy based analysis and modularity to be applied to such models.

### 3.1 Generating a bond graph from a stoichiometric matrix

A bond graph can be constructed from a stoichiometric matrix by using the following procedure:

1. For each *species* create a **Ce** component with appropriate name and a **0** junction; connect a bond from the **0** junction to the **Ce** component.
2. For each *reaction* create an **Re** component with appropriate name and two **1** junctions; connect a bond from one **1** junction to the forward port of the **Re** component and a bond from the reverse port of the **Re** component to the other **1** junction.
3. For each *negative* entry *N*_*ij*_ in the stoichiometric matrix, connect −*N*_*ij*_ bonds from the zero junction connected to the *i*th species to the the **1** junction connected to the forward port of the *j*th reaction.
4. For each *positive* entry *N*_*ij*_ in the stoichiometric matrix, connect *N*_*ij*_ bonds from the one junction connected to the reverse port of the *j*th reaction to the zero junction connected to the *i*th species.
5. If an Michaelis-Menten formulation is required, each **Re** component is replaced by a bond graph module (§ 3.2) corresponding to the enzyme catalysed reaction pair (8) and Appendix E.

For example, the reaction 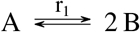 has the stoichiometric matrix

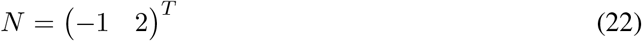

and the bond graph of Figure 1(a). The reaction 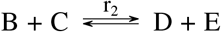 has the stoichiometric matrix

**Figure 1:**
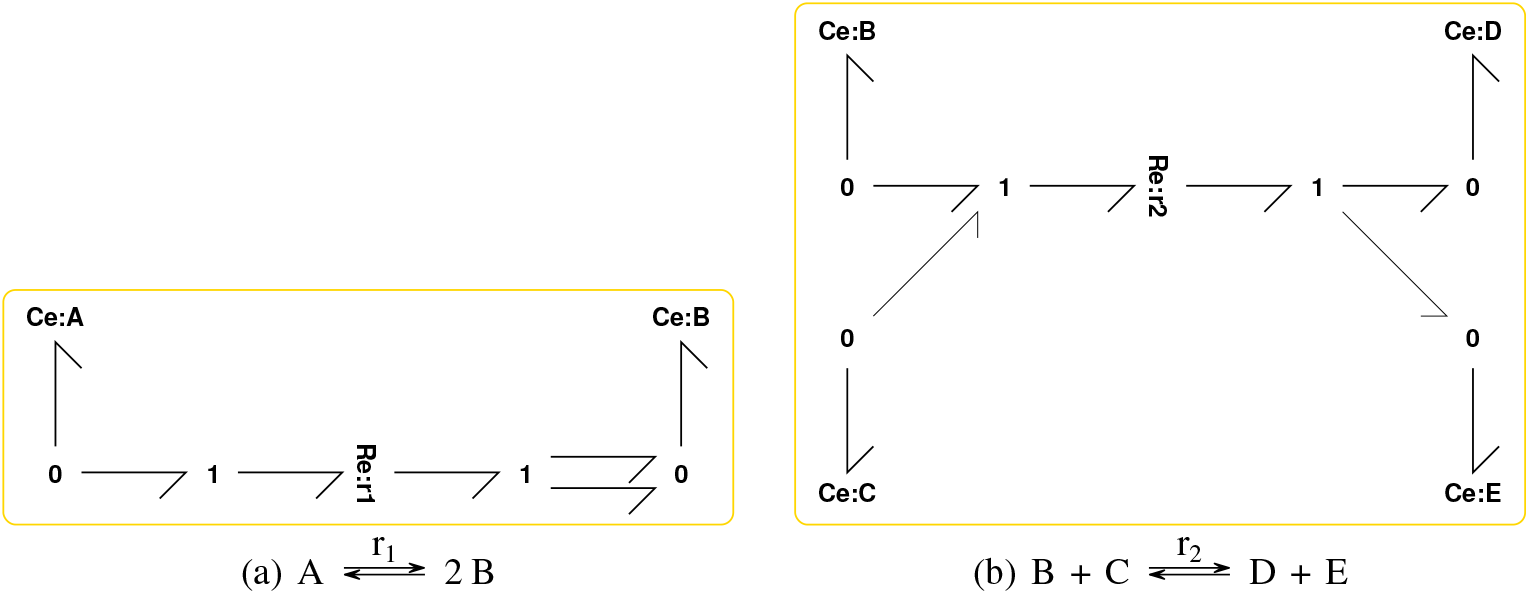
Bond graphs of simple reactions. (a) and (b) are used as modules M1 and M2 in § 3.2.

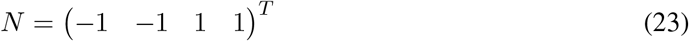

and has the bond graph of Figure 1(b).

Bond graphs provide a graphical representation of a system. While this provides an intuitive and clear visual representation when dealing with small systems such as the ones shown above, such visualisation becomes cumbersome for large systems. We employ two approaches to overcome this issue for the large-scale systems considered in this paper: modularity and a non-graphical (or programmatic) representation. In particular, we use a recent concept of bond graph modularity [38] in § 3.2 and the recently developed BondGraphTools package[63] (https://pypi.org/project/BondGraphTools/) as a non-graphical representation that allows large-scale systems to be constructed in a scalable and automated manner. This is discussed further below.

### 3.2 Modularity

Two related but distinct concepts of modularity [44] are *computational modularity*, where physical correctness is retained, and *behavioural modularity*, where module behaviour (such as ultra-sensitivity) is retained. Here we discuss computational modularity. In particular, it is shown how the concept of external flows, as discussed in § 2.2, is key to bond graph modularity.

Modular bond graphs provide a way of decomposing complex biomolecular systems into manageable subsystems [43, 44, 53]. This paper combines the modularity concepts of Neal et. al. [64–66] with the bond graph approach to give a more flexible approach to modularity. The basic idea is simple [38]. Modules are self-contained and have no explicit ports, but any species represented by a **Ce** component has the potential to become a port available for external connection. Thus, if two modules share the same species, the corresponding **Ce** component in each module is replaced by a port (labelled with the same name), and the species is explicitly represented as a **Ce** component in the parent model. This approach allows each module to be individually tested prior to being integrated into a larger model.

We use the following algorithm to merge bond graph models of stoichiometric networks:

1. Within each module, each **Ce** component corresponding to a common species is *exposed*, that is, replaced by a *port*, or external connection.
2. For each common species, create a **Ce** component connected to a **0** component.
3. Connect all module ports associated with each species to the **0** junction associated with the species; all instances of **Ce** components corresponding to each species are thus *unified* into the same component.

For example, let modules M1 and M2 correspond to Figures 1(a) & 1(b) respectively. The com-position of these modules requires the common species B to be exposed in both modules. This is illustrated in Figure2, where both modules are connected to the new **Ce**:**B** component via a **0** junction. The composite system contains the two coupled reactions:

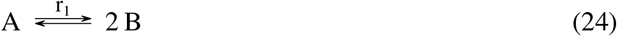

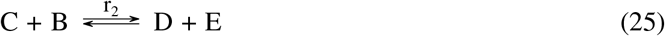

§ 4 gives examples of modular decomposition of a metabolic system and § 4.2 gives an example of how such modules can be combined using the methods of this section. The pathway analysis of § 2.2 can be applied to modules themselves, and to systems built of modules, to give insight into the overall behaviour of complex systems; this is illustrated in § 4.2.

**Figure 2:**
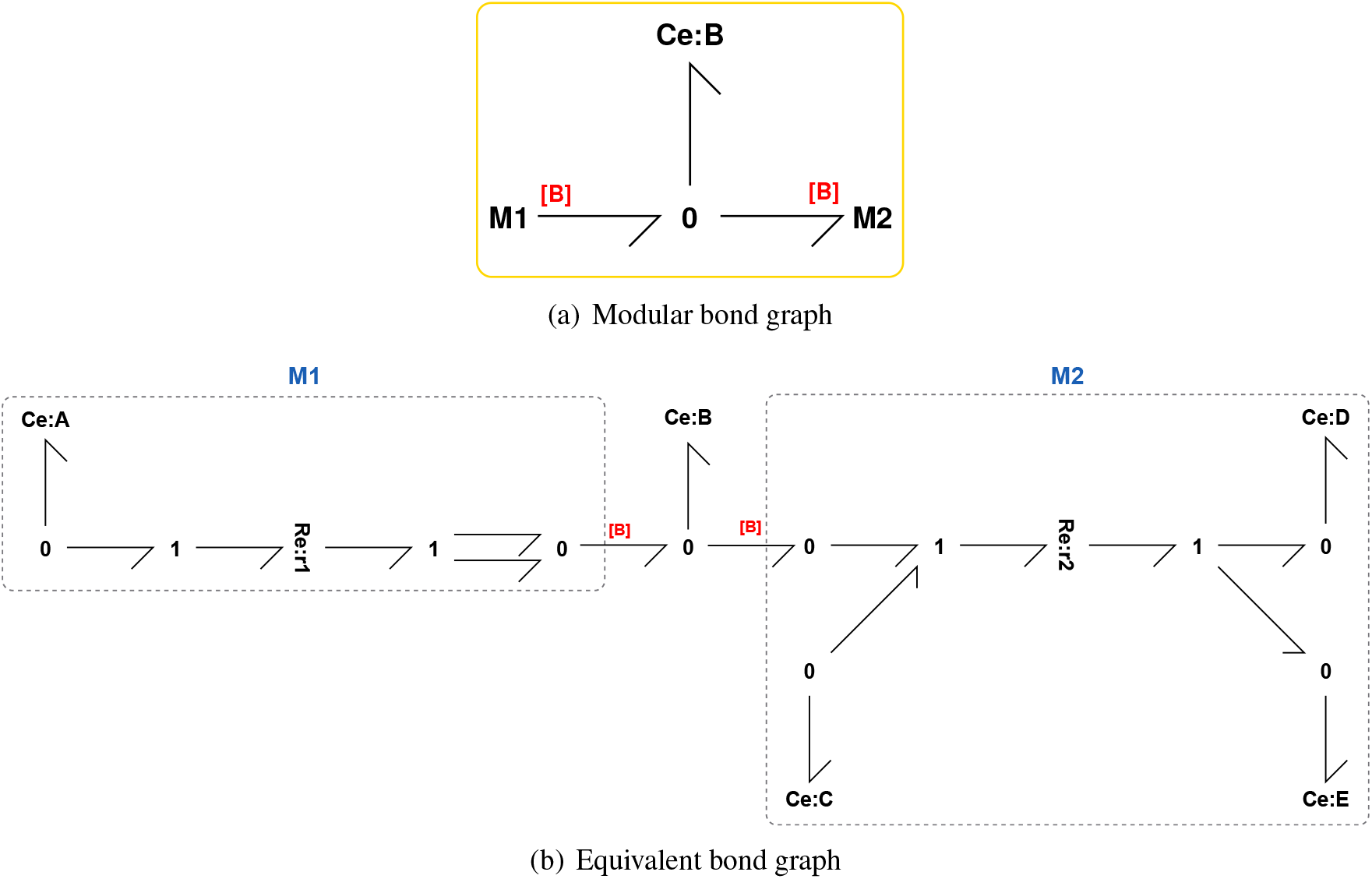
Modularity. Modules M1 and M2 correspond to Figures 1(a) & 1(b) respectively. The common species B is exposed as a port in each module and connected to the new **Ce**:**B** component via a **0** junction. (a) shows the compact modular form and (b) contains equivalent bond graph when the contents of the modules are expanded.

**Figure 3:**
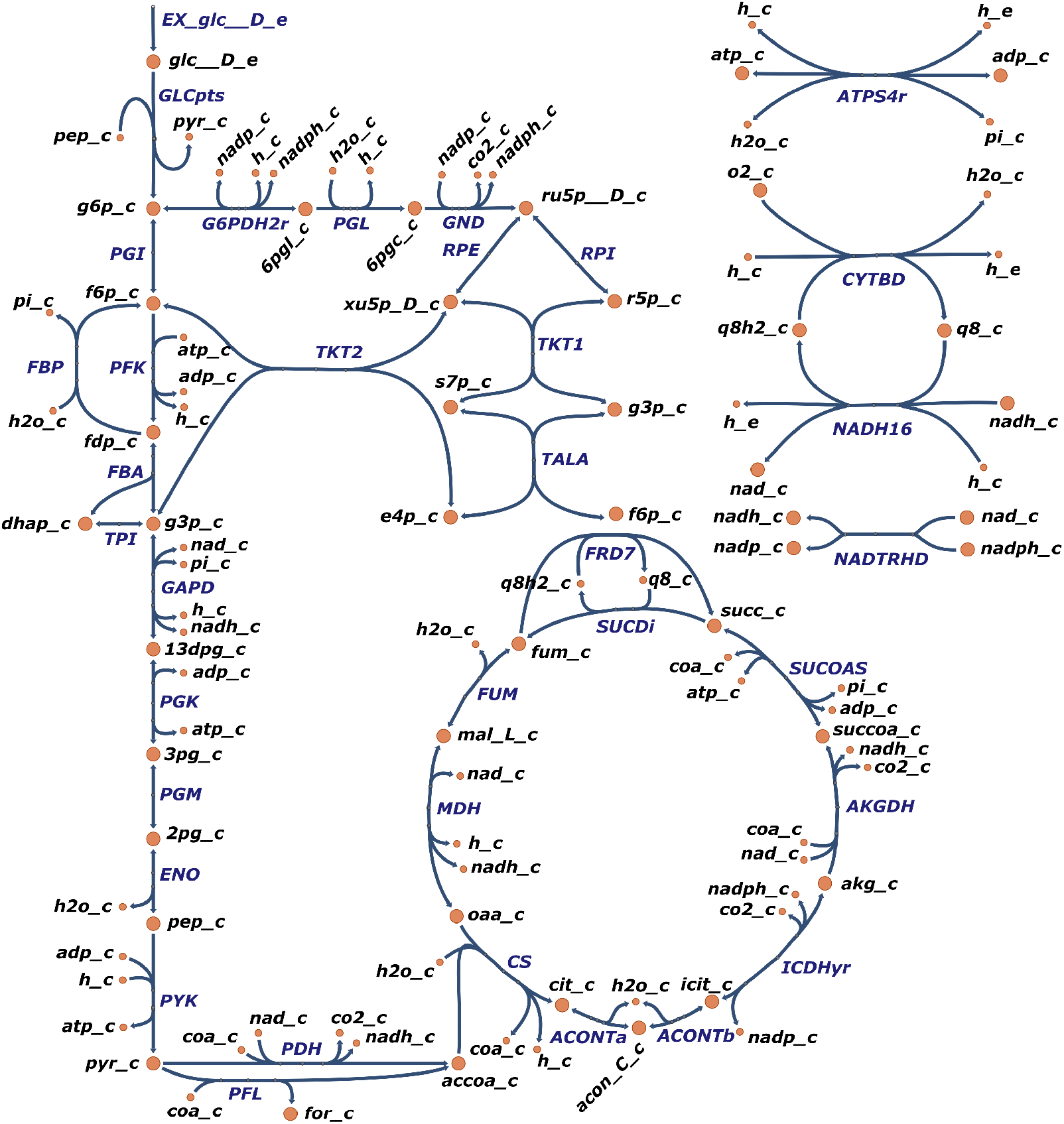
*E. coli* Core Model. The extracted reactions corresponding to the Glycolysis, Pentose-Phosphate Pathways and TCA cycle parts of the model are shown; a complete list of reactions is given in Appendix D. The diagram was created using Escher [68].

The concept of modularity can be extended to include common **Re** (reaction) components [67]; but this concept is not pursued in this paper.

### 3.3 Energy Balance Analysis (EBA) in a bond graph context

FBA [13] uses the linear Equation (19) within a constrained linear optimisation to compute pathway flows. EBA [16] adds two sorts of nonlinear constraint arising from thermodynamics. This section shows that the bond graph approach automatically includes the EBA constraint equations by considering Inequality (11) and Equation (18). In particular:

1. Inequality (11) corresponds to Equation 8 of Beard et al. [16]. This inequality can be reexpressed as:

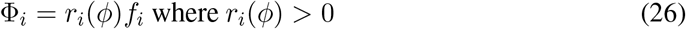

*r*_*i*_ corresponds to the “flux resistances” on p.83 of Beard and Qian [16].
2. If *K* is the right nullspace matrix of *N*, it follows from Equation (18) that

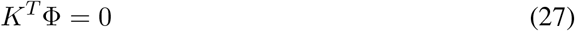 This corresponds to Equation 7 of Beard et al [16]. Note that *K* defines the pathways of the closed system system, with no chemostats.

Moreover, the pathways of the open system as defined by *K*^*cd*^ can be considered by defining *R* = diag *r*_*i*_ and using Equation (19):

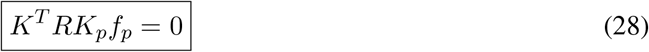

Equation (28) and inequality (26) constrain the pathway flows *f*_*p*_. This is illustrated in Appendix A.

## 4 Application to the *E. coli* Core Model

The *E. coli* Core Model [8, 51] is a well-documented and readily-available stoichiometric model of a biomolecular system; species, reactions and stoichiometric matrix were extracted from the CobraPy model: “textbook”. Using the methods of § 3.1, the corresponding bond graph model was created which, as discussed in the Introduction, automatically satisfies thermodynamic constraints.

To illustrate the concepts developed above, we analyse two subsets of reactions within this model:

1. § 4.1 uses the methods of § 2.2 to examine possible pathways within the system formed from the combined Glycolysis & Pentose Phosphate pathway (which produces precursors to the synthesis of nucleotides).
2. § 4.2 uses the modularity approach of § 3.2 to build a modular model of respiration using Glycolysis, the TCA cycle, the Electron Transport Chain and ATP synthase as modules. Furthermore, the methods of § 2.2 are applied to examine the pathway properties of an individual module (the TCA cycle) as well as the overall system.

### 4.1 Glycolysis & Pentose Phosphate pathway

The combination of the Glycolysis & Pentose Phosphate networks provides a number of different products from the metabolism of glucose. This flexibility is adopted by proliferating cells, such as those associated with cancer, to adapt to changing requirements of biomass and energy production [69, 70].

We construct a stoichiometric model of these pathways, consisting of the upper reactions of glycolysis and the pentose phosphate pathway. The full reaction network is given in Appendix C, and a bond graph is constructed using the methods of § 3.1.

As discussed in the textbooks [61, 71], it is illuminating to pick out individual paths through the network to see how these may be utilised to provide a variety of products. This is reproduced here by choosing appropriate chemostats and flowstats (§ 2.2) to give the results listed by Garrett and Grisham [61] § 22.6d. In each case, the corresponding pathway reaction potential is given. For consistency with Garrett and Grisham [61] § 22.6d, each pathway starts with Glucose 6-phosphate (G_6_P).

We use the following list of chemostats (together with additional chemostats) for the pathway analysis below: {ADP, ATP, CO_2_, G_6_P, H, H_2_O, NAD, NADH, NADP, NADPH, PI, PYR}. The pathways are generated using the methods of § 2.2.

1. R_5_P & NADPH generation **Chemostats:** RP5 **Flowstats:** PGI, TKT2 **Pathway: G6PDH2R + PGL + GND + RPI** **Reaction:** G_6_P + H_2_O + 2 NADP ⇄ CO_2_ + 2 H + 2 NADPH + R_5_P
2. R_5_P generation **Chemostats:** RP5 **Flowstats:** GAPD, G6PDH2R **Pathway: -5 PGI -PFK -FBA -TPI -4 RPI + 2 TKT2 + 2 TALA + 2 TKT1 + 4 RPE** **Reaction:** ADP + H + 6 R_5_P ⇄ ATP + 5 G_6_P
3. NADPH generation **Chemostats:** None **Flowstats:** GAPD **Pathway: -5 PGI -PFK -FBA -TPI + 6 G6PDH2R + 6 PGL + 6 GND + 2 RPI + 2 TKT2 + 2 TALA + 2 TKT1 + 4RPE** **Reaction:** ADP + G_6_P + 6 H_2_O + 12 NADP ⇄ ATP + 6 CO_2_ + 11 H + 12 NADPH

In § 5, we use the model of the Glycolysis & Pentose Phosphate pathways as a basis for inferring parameters from experimental data. Once the parameters have been identified (§ 5.4), dynamic simulations of these pathways can be run. This is shown later in § 5.6.

### 4.2 Respiration

To illustrate the utility of using bond graphs for the modular construction of stoichiometric models, we construct a model of respiration by combining the subsystems of Glycolysis, TCA cycle, Electron Transport Chain and ATP Synthase. Reactions for each of these subnetworks were extracted from the CobraPy model; these reactions are listed in Appendix D. For simplicity, reactions PDH and PFL (converting PYR to ACCOA) and reaction NADTRHD (converting NADP/NADPH to NAD/NADH) were included in the TCA cycle module. Once these are converted into bond graphs, the algorithm in § 3.2 was used to combine these models together into a model of respiration.

#### 4.2.1 Analysis of individual modules

An advantage of considering subsystems as separate modules is that these modules can be analysed individually. For example, the TCA cycle module can be analysed using the set of chemostats (see § 2.2):

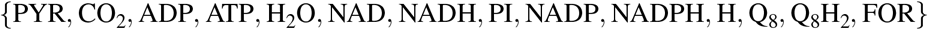

Three pathways result from this analysis:

1. **FRD7 + SUCDI** This is a type III pathway with no overall reaction.
2. **CS + ACONTA + ACONTB + ICDHYR + AKGDH + SUCOAS + FRD7 + FUM + MDH + PDH** This is a type I pathway with the reaction

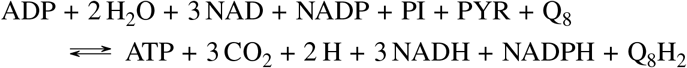
3. **CS + ACONTA + ACONTB + ICDHYR + AKGDH + SUCOAS + FRD7 + FUM + MDH + PFL** This is a type I pathway with the reaction

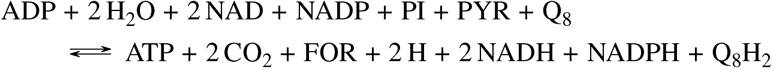

Pathways 2 and 3 utilise the potential of PYR to generate NADH, NADHP, ATP and Q_8_H_2_ whilst releasing CO_2_ and H.

#### 4.2.2 Analysis of combined network

The bond graph approach provides a method for easily combining stoichiometric models using the methods of § 3.2. Here we demonstrate this by constructing a model of respiration from the individual modules Glycolysis, TCA cycle, Electron Transport Chain and ATP Synthase. We begin by first combining the Glycolysis and TCA modules, as indicated in Figure 4(a). As well as the common species PYR (pyruvate) explicitly shown, the set of species

**Figure 4:**
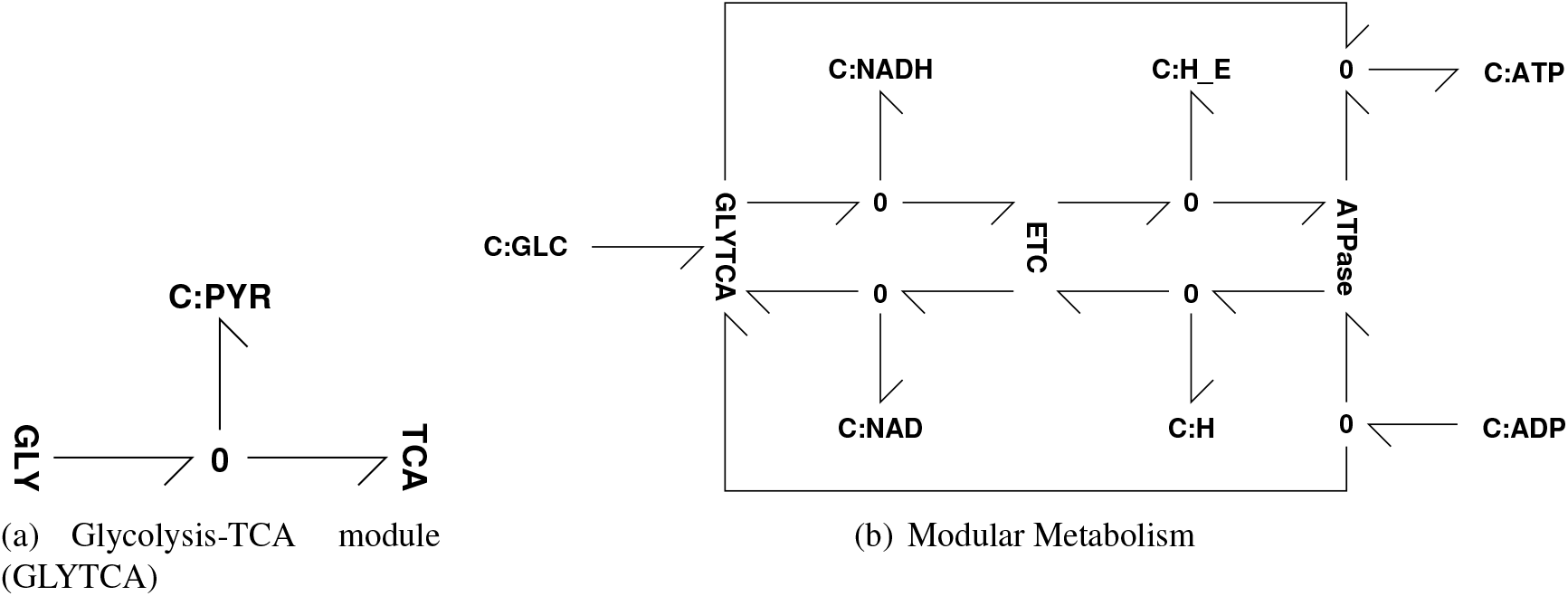
Modularity. (a) The two modules **GLY** (Glycolysis) and **TCA** (TCA cycle) each contain a bond graph representation of the relevant reactions. As discussed in § 3.2, they are combined into a single module by combining common species; in this case PYR is shown explicitly – other common species are {ATP, ADP, PI, H, NAD, NADH, H_2_O}. (b) The three modules **GLYTCA** (containing the two modules **GLY** and **TCA**), **ETC** and **ATP Synthase** are combined by unifying common species. This is shown for principle common species and emphasises that **ETC** is powered by NADH from **GLYTCA, ATP Synthase** is powered by the external protons H_E_ and both **GLYTCA** and **ATP Synthase** generate ATP from ADP. Common species not explicitly shown are *{* PI, H_2_O, Q_8_, Q_8_H_2_*}*.

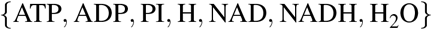

were also declared to be common.

The full model of respiration is then constructed by combining the Glycolysis+TCA cycle module with the Electron Transport Chain and ATP Synthase modules, as indicated in Figure 4(b). In addition to the common species explicitly shown

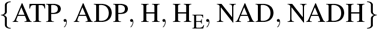

the set of species

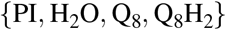

were also declared to be common.

To analyse this overall module, the chemostats were chosen to be:

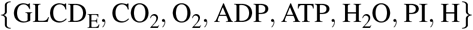

Using the methods of § 2.2, the three pathways in this network are

1. **PFK + FBP** This is a type II pathway with the overall reaction

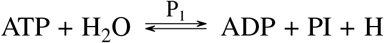

This Futile Cycle has regulatory implications[62].
2. **-FRD7 + SUCDI** This is a type III pathway with no overall reaction.
3. **2 GLCPTS + 2 PGI + 2 PFK + 2 FBA + 2 TPI + 4 GAPD + 4 PGK -4 PGM + 4 ENO + 2 PYK + 4 PDH + 4 CS + 4 ACONTA + 4 ACONTB + 4 ICDHYR + 4 AKGDH + 4 SUCOAS + 4 FRD7 + 4 FUM + 4 MDH + 4 NADTRHD + 20 NADH16 + 12 CYTBD + 27 ATPS4R**

This is a type I pathway with the reaction

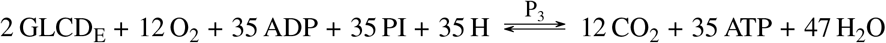

Pathway 3 corresponds to the metabolic generation of ATP using the free energy of GLCD_E_. The ratio of ATP to GLCD_E_ is 17.5; this is the value quoted by Palsson [8] § 19.2.

## 5 Dynamic Modelling and Parameter Estimation

Dynamic models of biochemical networks have the potential to aid the understanding of how sub-processes change over time, and can potentially elucidate important control structures within these networks [72]. However, due to their nonlinear nature, parameter estimation is one of the most challenging aspects of developing models of biomolecular systems [73].

Parameter estimation depends on both the form of the model and the type of data available. This section assumes a bond graph model with the mass-action kinetics of Equation (6) and that the following data are available for a single steady-state condition:

1. Reaction potentials Φ (equivalent to reaction Gibbs free energy).
2. Reaction flows *f*.
3. Species concentration *c*.

If data at three or more steady-state conditions were available, more complex kinetics such as the reversible Michaelis-Menten formulation (7) could be used but this is not pursued in this paper.

In recent times, such data are becoming more readily available; species concentrations can be obtained from metabolomics data, and tracer experiments involving ^13^C and ^2^H have been used to infer both fluxomics data for reaction flows [74, 75] and thermodynamic data for reaction potentials [74, 76, 77]. In the following examples, we make use of the dataset obtained by Park et al. [74] to infer the thermodynamic parameters using a relatively fast quadratic programming (QP) algorithm.

Because bond graph models are thermodynamically consistent, the estimated parameters have physical meaning and the resultant estimated model, though not necessarily correct, is physically plausible [47]. Moreover, physical constraints imply parametric constraints thus reducing the parameter search space.

### 5.1 Species potentials

Because of the energetic constraints implied by the bond graph, the reaction potentials Φ are related to the species potentials *ϕ* by Equation (13). Since some reaction potentials may be unavailable, we rearrange and partition Φ and the stoichiometric matrix *N* so that

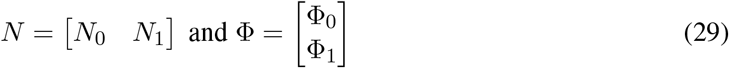

where Φ_0_ and Φ_1_ contain the known and unknown values of Φ respectively.

Given the measured value of Φ_0_ and the estimated species potentials 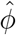, the *estimation error ϵ* is defined as:

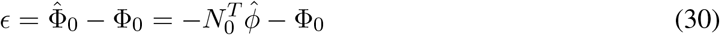

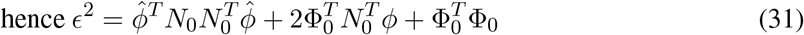

where the hat notation denotes estimated quantities. Although Φ_1_ is unknown, it is subject to the physical inequality (11). In this case, all of the measured flows are positive, hence inequality (11) can be combined with Equation (13) and rewritten as:

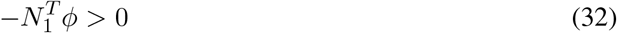

Equation (30) and inequality (32) can be embedded in a *quadratic program* (QP) [78]:

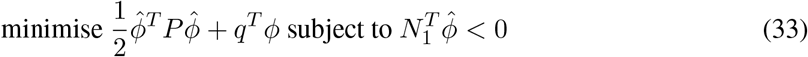

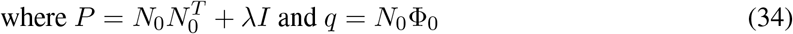

*I* is the *n*_*ϕ*_ *× n*_*ϕ*_ unit matrix and *λ >* 0 a small positive number. In some cases, there are more species than reactions and so the stoichiometric matrix *N* has more rows than columns. As a result, the number of species potentials *ϕ* is greater than the number of reaction potentials Φ and so Equation (13) has no unique solution for *ϕ* given Φ. Thus we use the *λI* term to which is required to turn a non-unique solution for *ϕ* into a minimum norm solution.

Having deduced a set of estimated species potentials 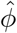 using the QP, the corresponding reaction potentials 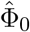 and 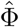 can be obtained from Equation (13) rewritten as:

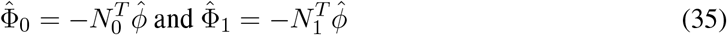

Once again, 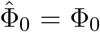 and the other values of Φ can be deduced from (35); because of the inequality constraint in the QP, these values are positive and thus physically plausible.

QP also handles equality constraints [78]; this provides a potential mechanism for incorporating known parameters into the procedure.

### 5.2 Pathway flows

From basic stoichiometric analysis, steady-state flows *f* can be written in terms of the pathway matrix *K*_*p*_ and pathway flows *f*_*p*_ by Equation (19) repeated here as

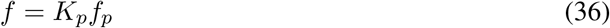

Note that, as discussed in § 2.2, the *pathway matrix K*_*p*_ is dependent on the choice of chemostats. In general, *K*_*p*_ has more rows than columns and thus the pathway flow *f*_*p*_ is over-determined by the reaction flows *f*. Hence, given a set of experimental flows *f*, an estimate 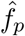 of *f*_*p*_ can be obtained from the *least-squares* formula:

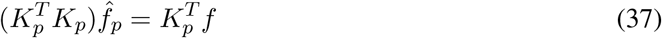

Note that:

1. 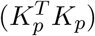 is a square *n*_*p*_ *× n*_*p*_ matrix where *n*_*p*_ is the number of pathways.
2. If some flows are not measured, the corresponding rows of *K*_*p*_ are deleted.
3. The reaction flows (including the missing ones) can be estimated from 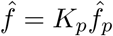.
4. From Equation (12), the estimated chemostat flows are given by the non-zero elements of

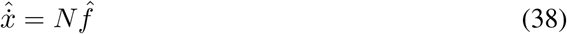

### 5.3 Reaction constants

In terms of estimated quantities, the reaction flow of Equation (6) can be rewritten as:

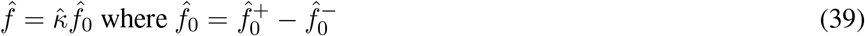

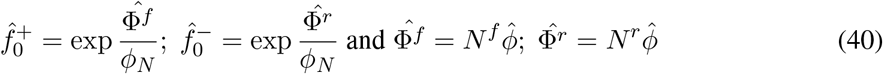

For each reaction, the estimated reaction constant 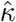 is then given by Equation (39).

Similarly, reversible Michaelis-Menten reaction kinetics can be written in terms of estimated quantities and three estimated parameters 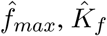 and 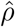 from Equation (7)

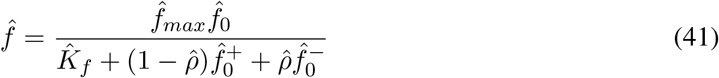

This can be rearranged as:

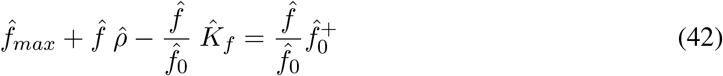

and can be in rewritten in linear-in-the-parameters form [79] as:

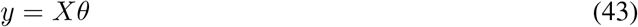

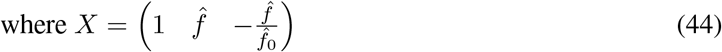

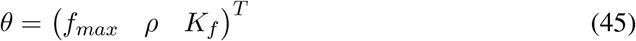

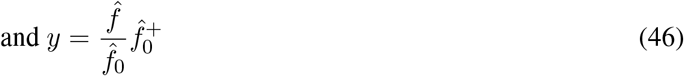

Given an estimate 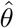 of *θ*, the *estimation error ϵ ′* is

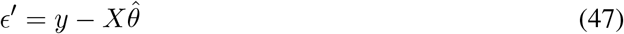

Because there are three unknown parameters (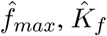 and 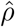), at least three different sets of steady-state data are required to uniquely determine the parameters; this case is not considered here. Alternatively, these unknown parameters can be determined using measured constants from the literature [29]. Such known parameters can be included using an *equality* constraint of the form 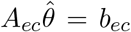 – an example appears in § 5.5. Noting that all elements of *θ* are positive, 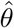 also has the *inequality constraint* 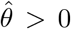, the error Equation (47) together with the constraints can be embedded *quadratic program* (QP) [78]:

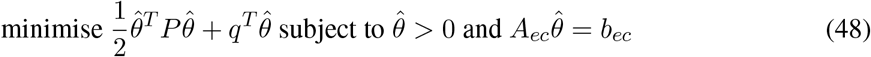

*I* is the *n*_*θ*_ *× n*_*θ*_ unit matrix and *λ >* 0 a small number. The parameters of the equivalent bond graph model can be deduced using Equation (9).

More general reaction kinetics [29] can be incorporated in a straightforward manner but, however, would require non-linear fitting procedures to determine parameters.

### 5.4 Dynamical parameters

The parameter *K* of the species components (**Ce**) determines the time course of species amounts and reaction flows when there is a deviation from steady-state. Using Equation (5), this can be determined from the species potential estimate 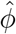 and the amount of species *x*^*⊖*^ at the steady-state conditions. Expressing amounts per unit volume, it follows that *x*^*⊖*^ = *c*, the species concentration at the steady-state conditions.

### 5.5 Parameters for the Glycolysis & Pentose Phosphate model

The bond graph of the Glycolysis & Pentose Phosphate model (§ 4.1) was parameterised to fit *E. coli* experimental data [74] using the approach described in this section. Table 2 [74] gives experimentally measured values of the reaction Gibbs energy Δ*G* for all of the reactions in the model except for G6PDH2R and PGL. The known values of Δ*G* were converted to reaction potentials Φ_0_ (mV). The unknown potentials Φ_1_ were constrained to be greater than 1 mV. The first column of Table 1(c) gives the experimental values of reaction potential Φ with the unknown values indicated by – the second column gives the corresponding estimates 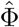 (mV). The estimated and known values are identical; of the two estimated unknown values, that for PGL lies on the constraint – unconstrained optimisation gives physically impossible negative value.

**Table 1:** Estimated flows and parameters; flows and concentration normalised by *f*_0_ and *c*_0_ (53). Missing data indicated by –. (a) Estimated species potentials 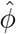 (§ 5.1), normalised concentration and species constants (§ 5.4). (b) Estimated chemostat flows (§ 5.2). (c) Estimated pathway flows (§ 5.2). (d) The estimated reaction potentials 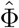; these are identical to the measured reaction potentials Φ where known (§ 5.1). The estimated reaction flows 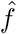 are close to the measured reaction flows where known (§ 5.2). The estimated mass-action reaction constants 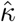 and the estimated Michaelis-Menten equivalent parameters 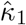 and 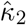 using (9) (§ 5.3) with *K*_*f*_ = 0.1 and *ρ* = 0.2.

| (a) Species parameters |  |  |  | (b) Chemostat flows |  | (c) Pathway flows |  |
| --- | --- | --- | --- | --- | --- | --- | --- |
| Species | $\hat{\phi}$ mV | $c/c_0$ | $\hat{K}$ | Chemostat | flow | Pathway | $\hat{f}_p$ |
| 6PGC | 29 | 0.4784 | 6.2335 | ADP | 63.12 | PPP1 | 63.12 |
| ADP | -27 | 0.0704 | 5.1546 | ATP | -63.12 | PPP2 | 7.98 |
| ATP | 27 | 1.2221 | 2.2539 | CO2 | 11.58 | PPP3 | 1.80 |
| CO2 | -30 | 0.0095 | 33.7942 | G3P | 128.03 |  |  |
| DHAP | -10 | 0.3883 | 1.7790 | G6P | -71.10 |  |  |
| E4P | -27 | 0.0062 | 57.9353 | H | 86.27 |  |  |
| F6P | -21 | 0.3198 | 1.4140 | H2O | -11.58 |  |  |
| FDP | -8 | 1.9289 | 0.3880 | NADP | -23.16 |  |  |
| G3P | -18 | 0.0344 | 14.9020 | NADPH | 23.16 |  |  |
| G6P | -5 | 1.0000 | 0.8377 | R5P | 6.19 |  |  |
| NADP | 30 | 0.0003 | 11747.0633 |  |  |  |  |
| NADPH | -30 | 0.0154 | 21.0027 |  |  |  |  |
| R5P | 5 | 0.0999 | 12.2419 |  |  |  |  |
| RU5PD | 5 | 0.0142 | 86.1551 |  |  |  |  |
| S7P | 24 | 0.1119 | 21.7513 |  |  |  |  |
| XU5PD | 5 | 0.0230 | 51.6829 |  |  |  |  |
| (d) Reaction flows and Parameters |  |  |  |  |  |  |  |
| Reaction | $\Phi$ mV | $\hat{\Phi}$ mV | $f/f_0$ | $\hat{f}/f_0$ | $\hat{\kappa}$ | $\hat{\kappa}_1$ | $\hat{\kappa}_2$ |
| PGI | 16.48 | 16.48 | 60.00 | 59.52 | 154.39 | 66.44 | 16.61 |
| PFK | 68.82 | 68.82 | 62.62 | 63.12 | 54.85 | 30.59 | 7.65 |
| FBA | 20.00 | 20.00 | 63.43 | 63.12 | 160.08 | 61.59 | 15.40 |
| TPI | 7.98 | 7.98 | 62.82 | 63.12 | 353.93 | 133.64 | 33.41 |
| G6PDH2R | – | 82.84 | – | 11.58 | 4.67 | 5.14 | 1.28 |
| PGL | – | 1.00 | – | 11.58 | 291.64 | 171.06 | 42.77 |
| GND | 114.53 | 114.53 | 11.70 | 11.58 | 1.27 | 4.78 | 1.19 |
| RPI | 0.04 | 0.04 | 7.87 | 7.98 | 4206.98 | 2785.35 | 696.34 |
| TKT2 | 16.38 | 16.38 | 0.91 | 1.80 | 9.17 | 2.24 | 0.56 |
| TALA | 54.41 | 54.41 | – | 1.80 | 1.66 | 0.94 | 0.23 |
| TKT1 | 4.04 | 4.04 | 2.92 | 1.80 | 8.82 | 6.66 | 1.67 |
| RPE | 0.83 | 0.83 | 3.83 | 3.59 | 96.07 | 63.27 | 15.82 |

As discussed in § 2.2, pathways are determined by chemostats. In this case it was assumed that the set of chemostats was: {ADP, ATP, CO2, G3P, G6P, H, H2O, NADP, NADPH, R5P}. Using the methods of § 2.2, there were three pathways:

1. PGI + PFK + FBA + TPI
2. G6PDH2R + PGL + GND + RPI
3. 2 PGI + 2 G6PDH2R + 2 PGL + 2 GND + TKT2 + TALA + TKT1 + 2 RPE

with pathway matrix *K*_*p*_ given by

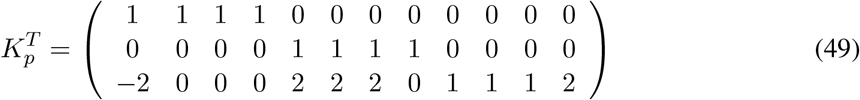

and corresponding reactions:

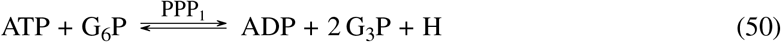

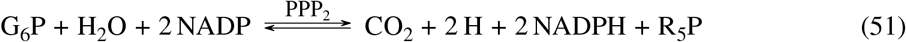

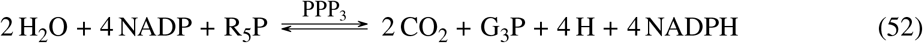

Data normalisation is important in the context of parameter identification in Systems Biology [80]. Here, the experimental concentration and flow data [74] was normalised with respect to the concentration of G_6_P and flow of PGI (given in mM/min) by defining:

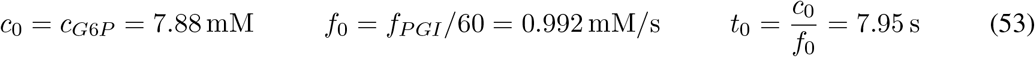

where *t*_0_ is the corresponding time unit.

Using the pathway decomposition and the method of § 5.2, the three pathway flows were deduced to be those of Table 1(d). The estimated reaction flows 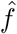 are then deduced from Equation (36) and given in the fifth column of Table 1(c). The chemostat flows are given in Table 1(b). The concentrations given in Table 3 [74] were used to derive the species parameters of Table 1(a).

The reaction constants *κ* of the mass action formulation are given in Table 1(d) together with the reaction constants *κ*_1_ and *κ*_2_ of the Michaelis-Menten formulation derived using the QP of (48). These parameters are used to perform a dynamical simulation in § 5.6.

### 5.6 Simulation

The parameters of Table 1(a)&(d) were used with the bond graph model of the Glycolysis & Pentose Phosphate pathway (§ 4.1) to run simulations. In § 4.1, we derived three pathways within this system; these are now simulated separately here. In particular, chemostats and flowstats (as defined in § 4.1) were implemented for the three cases and the initial concentrations were set to those in Table 1(a) where known and to unit values where unknown.

The simulation was performed separately for two cases: the mass-action formulation using the *κ* parameters and the Michaelis-Menten formulation using the 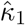 and 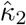 parameters.

Figure 5 shows the ratios 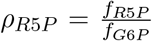 and 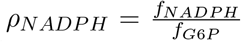 of the chemostat flows corresponding to the products R_5_P and NADH to the chemostat flow corresponding to the substrate G_6_P. At steady state, these ratios correspond to the stoichiometry of the three pathways of § 2.2. In particular, pathway i yields both products, pathway ii yields more R_5_P at the expense of NADPH and pathway iii yields more NADPH at the expense of R_5_P. Figures 5(a) and 5(b) correspond to the mass-action formulation and Figures 5(c) and 5(d) correspond to the Michaelis-Menten formulation.

**Figure 5:**
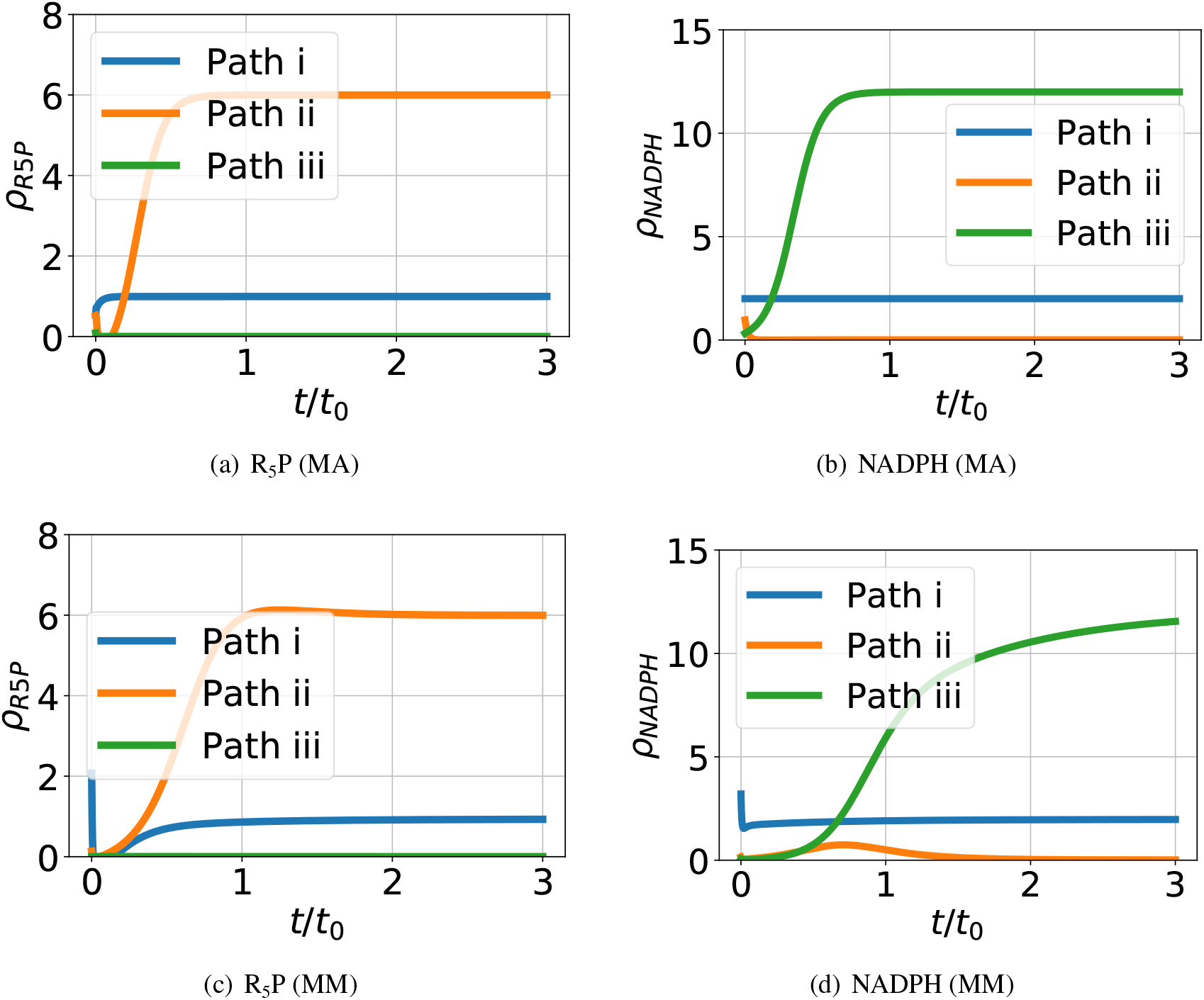
Pathway simulation. Ratios (*ρ*) of product (R_5_P & NADPH) chemostat flow to substrate (G_6_P) chemostat flow, plotted against time normalised by *t*_0_ (53), for each of the three pathways of § 2.2. The results are given for two cases: using the estimated mass-action (MA) parameter 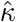 and using the estimated Michaelis-Menten equivalent parameters (MM) 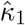 and 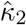 from Table 1. As discussed in § 2.2, pathway i yields both products, pathway ii yields more R_5_P at the expense of NADPH and pathway iii yields more NADPH at the expense of R_5_P. In this case both MA and MM give the same steady-state values but with differing dynamic response.

Because the two-reaction Michaelis-Menten formulation of enzyme catalysed reactions (8) explicitly includes the enzyme, such models can be used to examine system behaviour as enzyme levels change.

## 6 Conclusion

The formulation of dynamic simulation models for large-scale biological systems remains a key challenge in systems biology. With the advent of genome-scale simulation and whole-cell modelling, there is increasing recognition of the need for a modular approach in which model components can be formulated, tested and validated independently, and then seamlessly integrated together to form a model of the whole system. However, a dynamic modelling framework which is modular and which can in principle describe the broad range of biochemical and biophysical cellular processes has been elusive.

Several authors have acknowledged the need for energetic considerations to be integrated into modelling approaches, both to ensure that models are consistent with basic thermodynamic principles, and to enable calculation of energy flows and related concepts such as efficiency [67]. Here we have shown that thermodynamically compliant dynamic models of metabolism can be generated using the bond graph modelling approach, with the stoichiometric matrix as the starting point. Bond graphs, first advocated in the context of biological network thermodynamics by Oster et al. [34], represent both energy and mass flow through the biochemical network. Bond graphs separate the system connectivity from energy-dissipating processes (reactions), and thus are a very natural fit to network-based modelling in systems biology. As a port-based modelling approach, bond graphs are also inherently modular. Furthermore, application of bond graph modelling principles automatically endows models with a number of necessary features for large-scale modelling including modularity, thermodynamically distinguished parameters (wherein system-wide thermodynamic parameters relating to biochemical species are distinguished from reaction-specific parameters) and hence, as noted by Mason and Covert [29], improved opportunity for parameter identification from data.

Energy-based modelling of biochemical reaction networks using bond graphs naturally encompasses the EBA approach [16], where we have shown that the key equations of EBA are implicit in the system bond graph. This is a powerful advantage as it means that no additional steps are required in order to satisfy thermodynamic constraints. Any model formulated as a bond graph implicitly satisfies these constraints; it is not possible to impose, or infer from data, parameters which break these constraints.

A further benefit for large-scale modelling is that bond graphs naturally lend themselves to model reduction, for example through generation of reduced-order models using pathway analysis [53, 81]: any such simplified model will also satisfy the same thermodynamic constraints. This enables a hierarchical approach to modelling, and it is not necessary to model all aspects of the system at the same level of detail. Different levels of representation can be used as required, for example reflecting available knowledge and data about different parts of the system.

As noted in the Introduction, a key challenge in the development of dynamic models is the fitting of parameters to experimental data. We have shown that both mass-action kinetics and (reversible) Michaelis-Menten kinetics fall within the bond graph framework and therefore have a thermodynamically safe parameterisation; moreover, it is shown this parameterisation leads to a linear-in-the parameters estimation problem. Bond graphs separate the constitutive relations describing the reactions from the connectivity of the model; it is therefore possible to incorporate more complex kinetic schemescĩtepCor12, including inhibition, allosteric modulation and cooperativity within the bond graph approach thus retaining thermodynamically safe parameterisation. However, the resultant parameter estimation problem will not, in general, be linear-in-the parameters and will therefore require an optimisation approach such as that used by K-FIT [82]. Optimisation approaches such as K-FIT do not use a set of parameters that is thermodynamically safe by design, hence they need to derive additional constraints to incorporate thermodynamic consistency. Future work will examine how the thermodynamically safe parameterisation induced by the bond graph approach can be used to simplify such optimisation when applied to large systems and data sets.

According to Noor et al. [83] in the context of obtaining biological insights though omics data integration: “To maximize predictive power and mechanistic insights on the molecular level, ODE simulations based on physical models of binding and catalysis remain the gold standard.” The illustrative example of this paper shows how data involving flows, concentrations and chemical potentials can be integrated using the physical model structure provided by combining stoichiometric and bond graph approaches. It is believed that this provides a basis for integrating the larger and more varied omics data becoming available. Moreover, the physical basis of the approach can be used to indicate what additional data should be gathered to fully parameterise the model.

Here we have demonstrated that thermodynamically compliant dynamic models can be constructed starting from the stoichiometric matrix. The plethora of existing stoichiometric models for metabolic networks provides a natural starting point for this endeavour. However, while metabolic models are of central importance in a number of contexts, models of cellular physiology in general, and whole-cell models in particular, require a framework that can incorporate a much broader range of cellular processes, feedback and regulation. As a general tool for physically plausible systems modelling, bond graphs can naturally include energy compliant connections to other physical domains and processes, including transport [84], electrochemical transduction [38, 39], membrane potential dynamics [41], mechanochemical transduction and photosynthesis. Furthermore, through incorporation of control-theoretic concepts, enzyme modulation and feedback control can be represented in a coherent manner [62]. There remain however several key domains of cellular biology where to our knowledge there are as yet no examples of bond graph modelling, including transcription and translation [20, 85, 86]. These will need to be demonstrated in order to provide a complete road map for construction of modular and thermodynamically compliant whole-cell models using bond graphs.

## Data Accessibility

The figures and tables in this paper were generated using the Jupyter notebooks and Python code available at https://github.com/gawthrop/GawPanCra21.

## Acknowledgements

PJG would like to thank the Faculty of Engineering and Information Technology, University of Melbourne, for its support via a Professorial Fellowship. This research was in part conducted and funded by the Australian Research Council Centre of Excellence in Convergent Bio-Nano Science and Technology (project number CE140100036). The authors would like to thank the referees for their constructive comments.

## A EBA examples

These examples refer to § 3.3 and drawn from Beard et al [16].

### A.1 Example: Parallel reactions

Beard et al [16] motivate EBA using the example of two resistors in parallel. Figure 6(a) shows the bond graph of the analogous reaction system: the species A and B are joined by two reactions:

**Figure 6:**
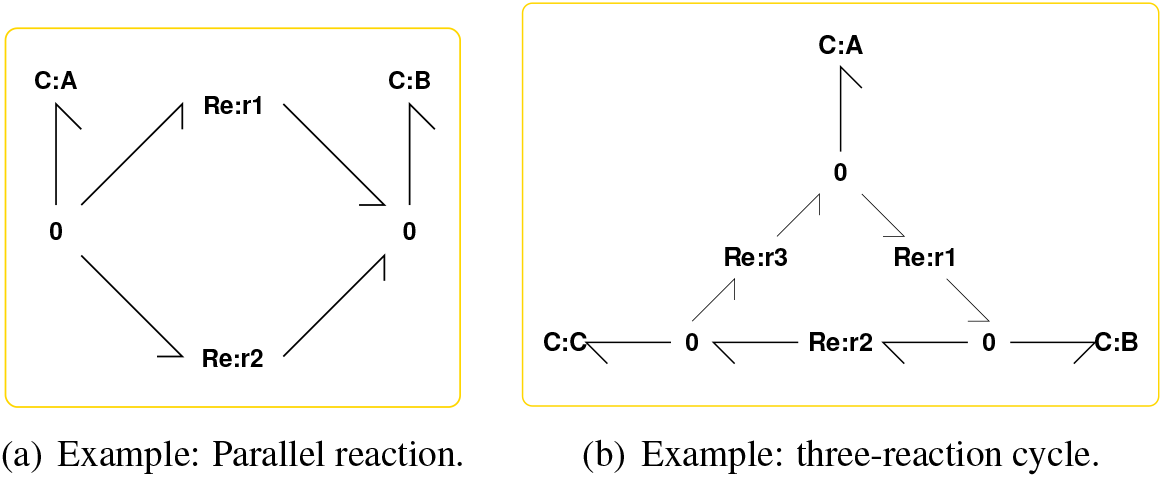
Bond graphs corresponding to examples from Beard et al [16] (**1** junctions are not shown for clarity)). (a) Beard et al [16, Fig. 2], (b) Beard et al [16, Fig. 3]

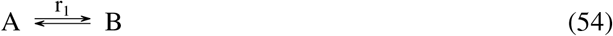

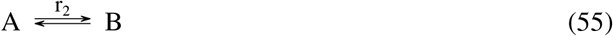

The stoichiometric matrix is:

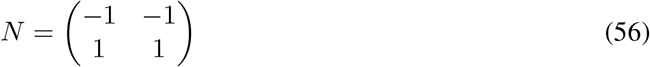

and the null space matrix *K* is

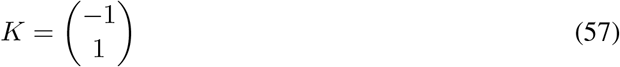

corresponding to the pathway: − *r*_1_ + *r*_2_.

Setting A and B as chemostats:

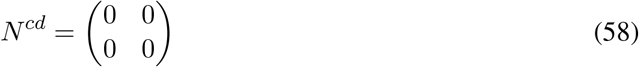

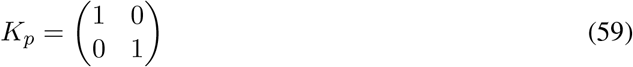

Equation (28) then becomes:

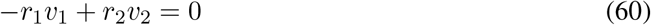

As *r*_*i*_ *>* 0, it follows that *v*_1_ and *v*_2_ must either be zero or have the same sign.

### A.2 Example: three-reaction cycle

Beard et al [16] give the example of a three-reaction cycle. Figure 6(b) shows the corresponding bond graph. The species A, B and C are joined by three reactions:

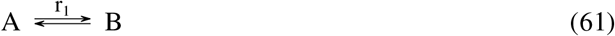

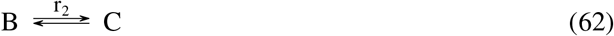

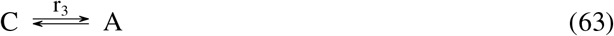

The stoichiometric matrix is:

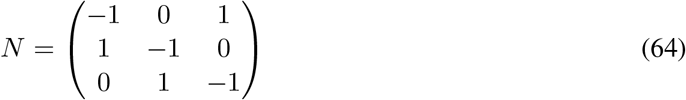

and the null space matrix *K* is

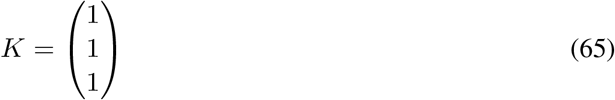

corresponding to the pathway: *r*_1_ + *r*_2_ + *r*_3_.

Setting A and B as chemostats:

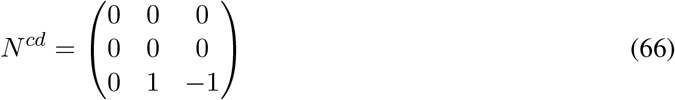

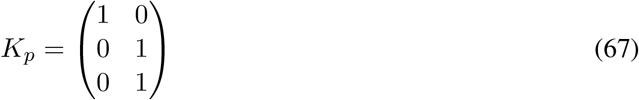

Equation (28) then becomes:

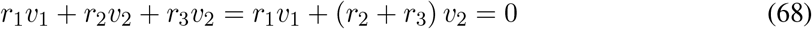

As *r*_*i*_ *>* 0, it follows that *v*_1_ and *v*_2_ must either be zero or have the opposite sign.

Alternatively, setting A, B and C as chemostats:

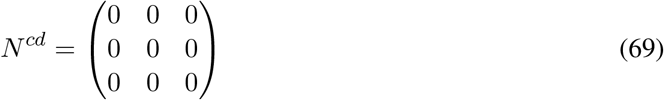

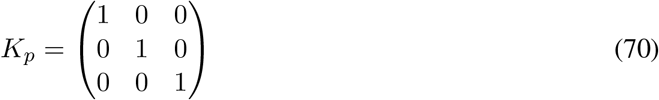

Equation (28) then becomes:

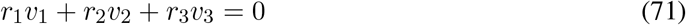

As *r*_*i*_ *>* 0, there are three possibilities: all flows are zero; one of the three pathway flows must have one sign and the other two flows the opposite sign; or one flow is zero and the other two have opposite signs.

## B Pathways: Illustrative example

This example refers to § 2.2. Noor [19] gives a simple illustrative example of the three types of pathway; Figure 7(a) gives the corresponding bond graph. the reactions are:

**Figure 7:**
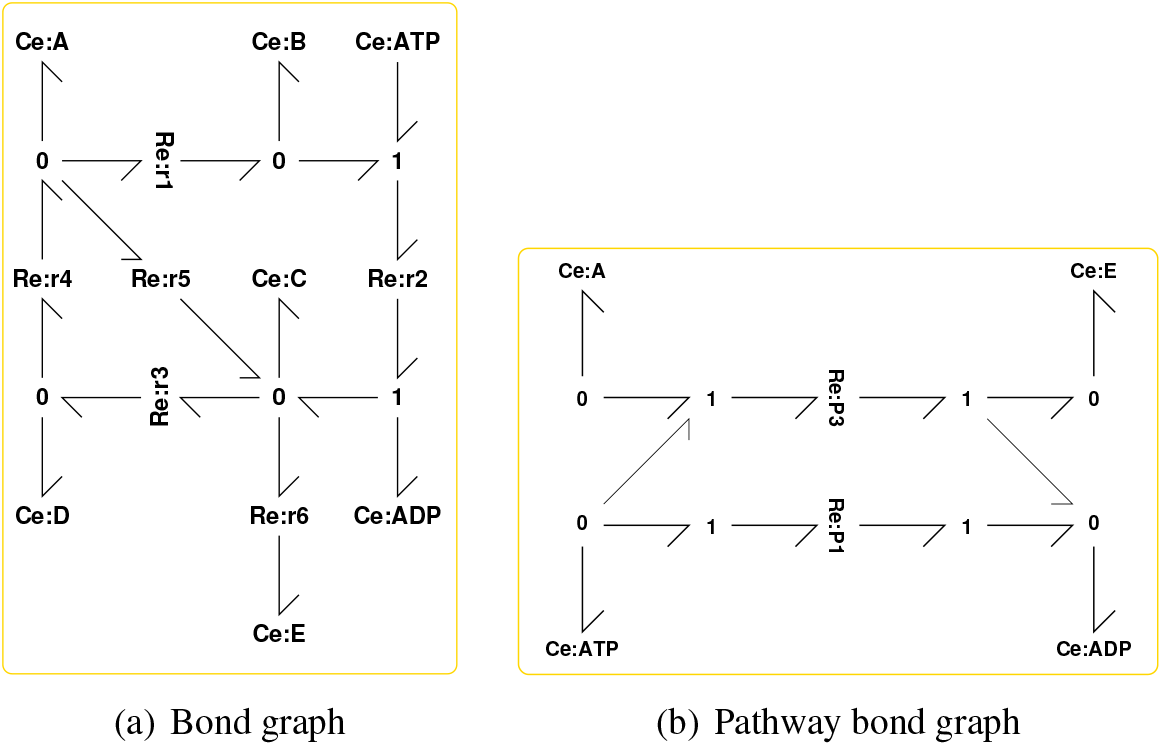
Bond graphs for illustrative example [19]

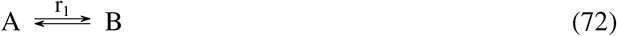

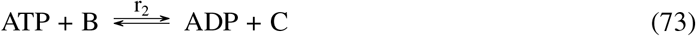

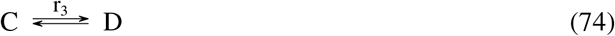

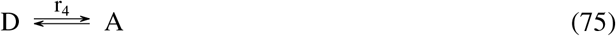

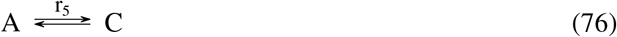

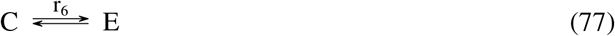

The there are seven species and six reactions giving states *x* and flows *v*:

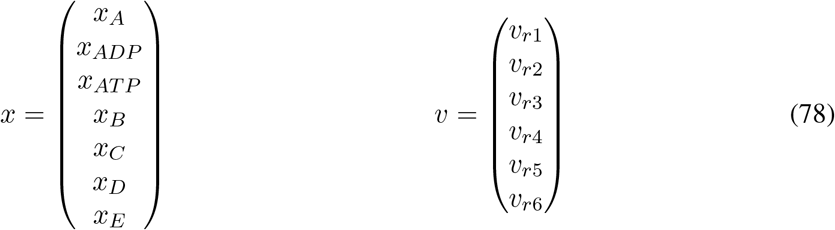

The stoichiometric matrix is:

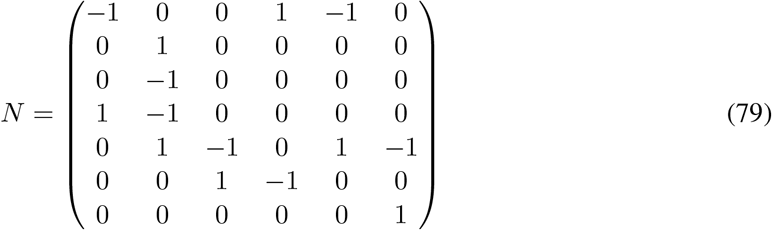

Setting A, E, ATP and ADP as chemostats, *N*^*cd*^ is constructed by setting the corresponding rows of *N* to zero. The corresponding null space is three dimensional and corresponds to the three pathways:

1. r1 + r2 + r3 + r4
2. r3 + r4 + r5
3. r1 + r2 + r6

Using (20), the pathway stoichiometric matrix *N*_*p*_ is:

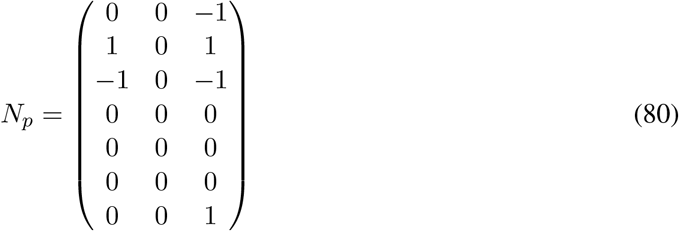

The three pathway reactions are:

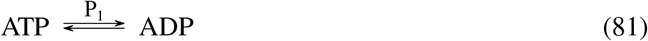

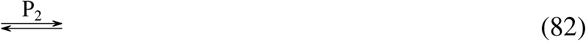

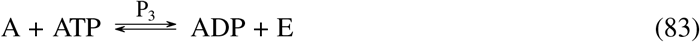

Pathway reaction P1 corresponds to a type II pathway, pathway reaction P2 to a type III pathway and pathway reaction P3 to a type I pathway where A is converted to E driven by the conversion of ATP to ADP. The example is extended by assigning a set of nominal chemical potentials 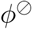 to the species: 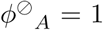, *ϕ*_*ATP*_ = 0, *ϕ*_*ADP*_ = 3, 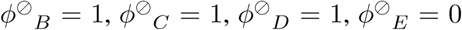. The pathway reaction potentials are then computed using (21) as Φ_*P*1_ = − 2, Φ_*P*2_ = 0, Φ_*P*3_ = − 1. As the potential for each pathway only depends on the species appearing in the pathway reactions, the potential of non-chemostatted species are irrelevant for this computation. In fact the potentials of the species will correspond to the steady-state values of concentrations of the non-chemostatted species arising from the flow patterns corresponding to the chemostat potentials [87]. The pathway bond graph appears in Figure 7(b).

## C Glycolysis & Pentose Phosphate Pathways

This section contains the reactions used in § 4.1 to generate the three pathways arising from the upper reactions of glycolysis and the pentose phosphate pathway. The reactions are extracted as discussed in § 4.

The reactions are:

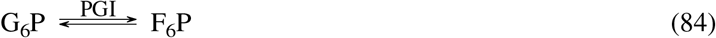

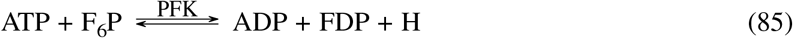

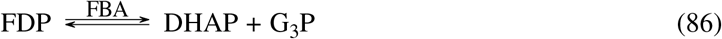

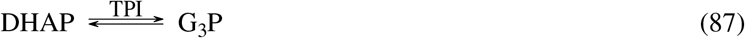

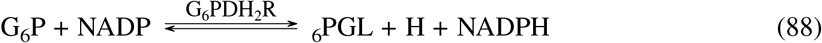

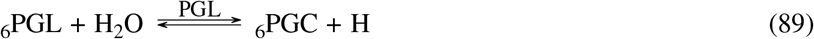

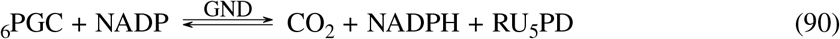

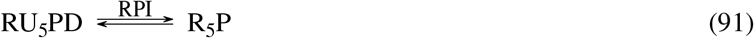

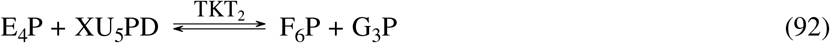

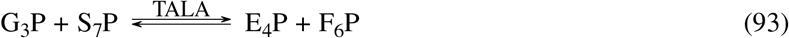

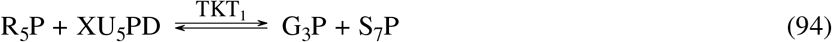

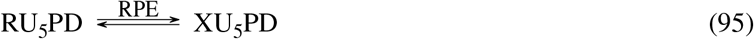

## D Modular representation of Metabolism: Reactions

This section contains the reactions used in § 4.2 which illustrates the utility of using bond graphs for the modular construction of stoichiometric models by constructing a model of respiration by combining the modular subsystems: Glycolysis, TCA cycle, Electron Transport Chain and ATP Synthase.

The reaction CYTBD (containing 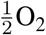) was multiplied by 2 to give integer stoichiometry and, for clarity, the reactions RPI, PGK, PGM, SUCOAS and FRD7 were reversed to give the conventional direction.

### D.1 Glycolysis

The reactions extracted are:

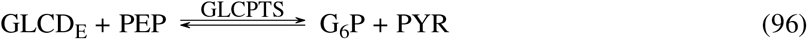

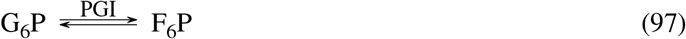

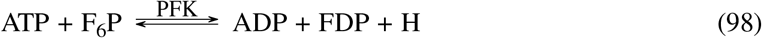

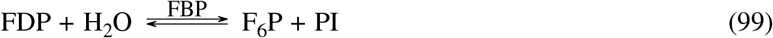

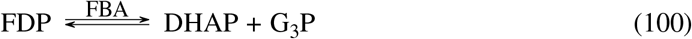

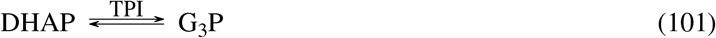

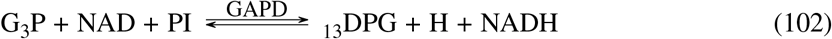

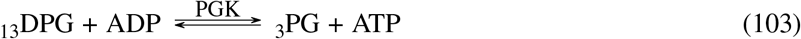

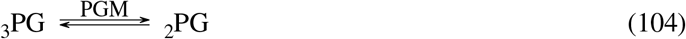

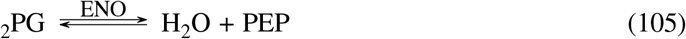

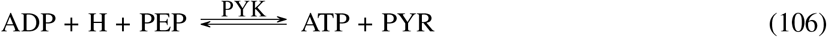

### D.2 TCA cycle

As well as the TCA cycle itself, this module includes:

1. the pyruvate (PYR) connection reactions: PDH and PFL and
2. the NAD/NADP interconversion reaction NADTRHD.

The reactions extracted are:

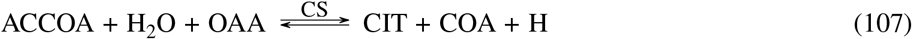

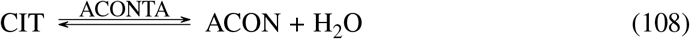

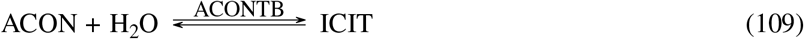

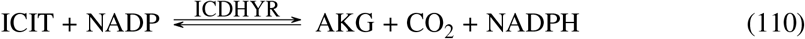

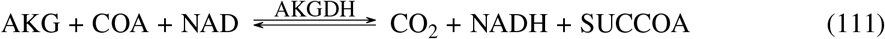

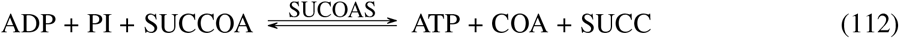

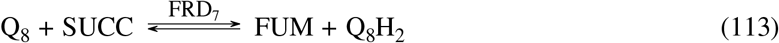

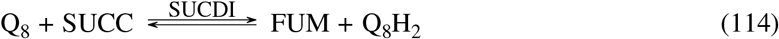

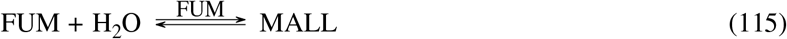

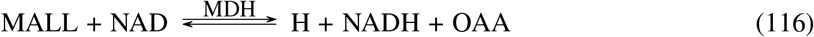

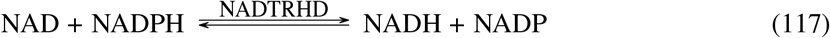

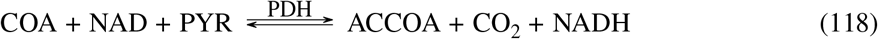

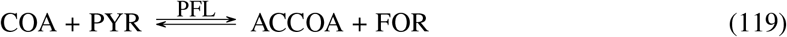

### D.3 Electron Transport Chain

The reactions extracted are:

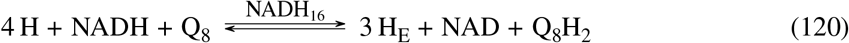

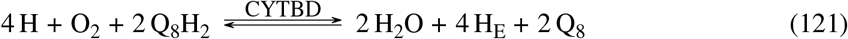

### D.4 ATP Synthase

The reaction extracted is:

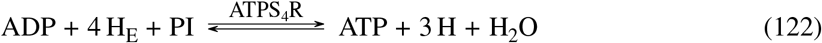

## E Enzyme-catalysed Reaction

**Figure 8:**
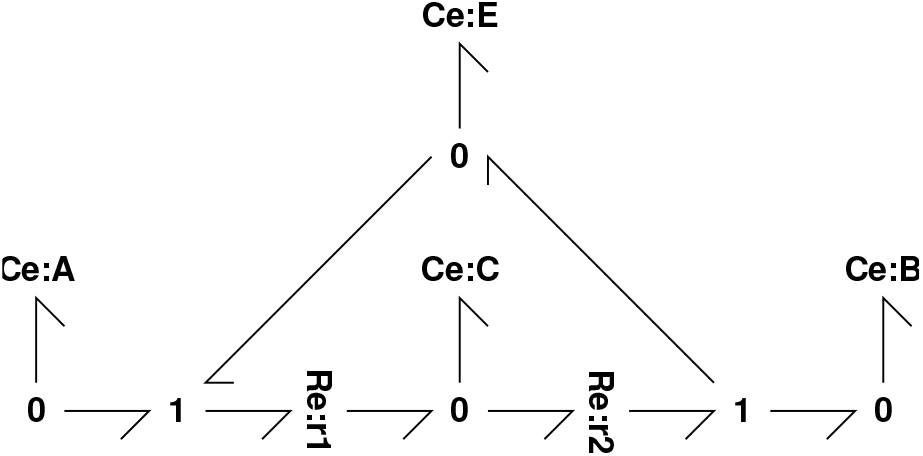
Bond graph representation of an Enzyme-catalysed Reaction

